# Protein-coding mutation in *Adcy3* increases adiposity and alters emotional behaviors sex-dependently in rats

**DOI:** 10.1101/2024.06.16.598846

**Authors:** Mackenzie Fitzpatrick, Alexandria Szalanczy, Angela Beeson, Anusha Vora, Christina Scott, Michael Grzybowski, Jason Klotz, Nataley Der, Rong Chen, Aron M. Geurts, Leah C. Solberg Woods

## Abstract

1

**Objective:** *Adenylate cyclase 3 (Adcy3)* has been linked to both obesity and major depressive disorder (MDD). Our lab identified a protein-coding variant in the 2^nd^ transmembrane (TM) helix of *Adcy3* in rats, and similar obesity variants have been identified in humans. The current study investigates the role of a TM variant in adiposity and behavior.

**Methods:** We used CRISPR-SpCas9 to mutate the TM domain of *Adcy3* in WKY rats (Adcy3^mut/mut^). We also created a heterozygous knockout rat in the same strain (Adcy3^+/-^). Wild-type (WT), Adcy3^+/-^, and Adcy3^mut/mut^ rats were fed a high-fat diet for 12 weeks. We measured body weight, fat mass, glucose tolerance, food intake, metabolism, emotion-like behaviors, and memory.

**Results:** Adcy3^+/-^ and Adcy3^mut/mut^ rats weighed more than WT rats due to increased fat mass. There were key sex differences: adiposity was driven by increased food intake in males but by decreased energy expenditure in females. Adcy3^mut/mut^ males displayed increased passive coping and decreased memory while Adcy3^mut/mut^ females displayed increased anxiety-like behavior.

**Conclusions:** These studies show that the ADCY3 TM domain plays a role in protein function, that *Adcy3* may contribute to the relationship between obesity and MDD, and that sex influences the relationships between *Adcy3*, metabolism, and behavior.

**Study Importance Questions:** *What is already known about this subject?:*

- *Adcy3* has been linked to both obesity and depression in humans, with protein-coding variants in the transmembrane domain leading to increased obesity. Yet the underlying mechanisms are unknown, and it is unknown if the same *Adcy3* variants can cause both obesity and depression.
- Rodent *Adcy3* knockout models show increased adiposity and altered emotional behaviors, but no rodent models have assessed the role of protein-coding variants in *Adcy3.* Very few studies have assessed sex differences.

*What are the new findings in your manuscript?:*

- This is the first study to assess the role of a protein-coding variant in the transmembrane domain of *Adcy3*, showing increases in both obesity and emotion-like behaviors in a rat model.
- We observed striking sex differences, where male and female rats had different factors driving their adiposity and different patterns of altered behavior.

*How might your results change the direction of research or the focus of clinical practice?:*

- Understanding the role of *Adcy3* in obesity and MDD may lead to improved treatments for both diseases.
- Improving our understanding of the sex differences caused by the same *Adcy3* mutation emphasizes the importance of studying both sexes and paves the way for personalized medicine approaches.

## 2 Introduction

Obesity rates continue to rise, and it has recently been predicted that the global prevalence of overweight and obesity will reach 51% by 2035.^1^ Similarly, global rates of major depressive disorder (MDD) have increased roughly 50% since 1990.^2^ While obesity and MDD have been shown to be bi-directionally associated,^3, 4^ the biological mechanisms underlying this association have not been fully elucidated. Nevertheless, genetic factors have been shown to play an important role in the development of both diseases.^5, 6^

*Adenylate cyclase 3* (*Adcy3*) has been established as an important gene for both obesity and MDD, supported by both human^7–21^ and rodent^22–28^ studies.^29^ ADCY3 is a plasma membrane-associated protein that catalyzes the synthesis of cAMP, a secondary messenger that has itself been linked to the development and treatment of obesity and MDD.^7, 8, 30, 31^ The causes of adiposity in *Adcy3* knockout (KO) models have not yet been firmly established. Some,^23, 24, 26^ but not all,^22, 25^ studies have reported that KO mice consume more food. Also, increased energy expenditure (EE) played a role in protecting mice with a gain-of-function mutation in *Adcy3* from diet-induced obesity.^27^ Increased depression- and anxiety-like behaviors and decreased memory have been reported in *Adcy3* KO mice,^25, 28^ although no studies have looked at behavioral and metabolic phenotypes in the same cohort. Furthermore, while most studies assessing *Adcy3* KO mice have only assessed males, one study that assessed females found that KO females required a high-fat diet to display the same traits as males.^22^ Behavior in female *Adcy3* KO mice has not yet been studied.

Using outbred rats, our lab identified a protein-coding variant in the 2^nd^ transmembrane (TM) helix of ADCY3 that is associated with adiposity,^32^ findings which have since been validated.^33, 34^ Similar TM protein-coding variants have been identified in humans in association with childhood obesity,^13, 17^ monogenic obesity,^14^ and severe adulthood obesity,^15^ emphasizing the translational importance of this region for metabolic health. TM domain mutations have not yet been specifically linked to MDD.

Critically, although both expression-altering and protein-coding variants in *Adcy3* have been identified in humans, only *Adcy3* KO models have been studied in rodents. No functional assessments of any *Adcy3* protein-coding variants have been conducted thus far even though most *Adcy3* variants identified in humans are not predicted to cause a full KO of this gene. Additionally, very few studies have assessed sex differences in the role of *Adcy3* in adiposity or behavior, particularly in the same rodent model. Thus, several gaps in knowledge still exist in our understanding of the role of *Adcy3* variants in obesity, in MDD, and in both diseases together. In the current study, we investigated how a protein-coding TM domain mutation in *Adcy3* affects adiposity, metabolism, and behavior in both sexes in an inbred rat model. We also investigated these same phenotypes in an *Adcy3* KO model to compare the effects of a KO vs. protein-coding mutation in this gene.

## 3 Methods

### 3.1 Animals

*Adcy3* knockout (KO) (WKY-*Adcy3*^em3Mcwi^; RRID:RGD_19165366) and Adcy3^mut/mut^ (WKY-*Adcy3*^em2Mcwi^; RRID:RGD_19165364) rats were generated by injecting recombinant SpCas9 protein (QB3 Macrolab, Berkeley, CA) along with a single guide RNA (IDT) targeting the *Adcy3* exon 2 sequence (CCTTTTTGCAGAGCACGAAA) into Wistar-Kyoto (WKY) rat embryos (WKY/NCrl; RRID:RGD_1358112). In *Adcy3* KO, a single base pair deletion caused a frameshift and whole-body KO of *Adcy3*. In Adcy3^mut/mut^, a 3 base pair deletion caused a F122delV123L mutation (**Figure 1A**). Heterozygous breeder pairs were maintained as described in Supplemental Methods.

**Figure 1.**
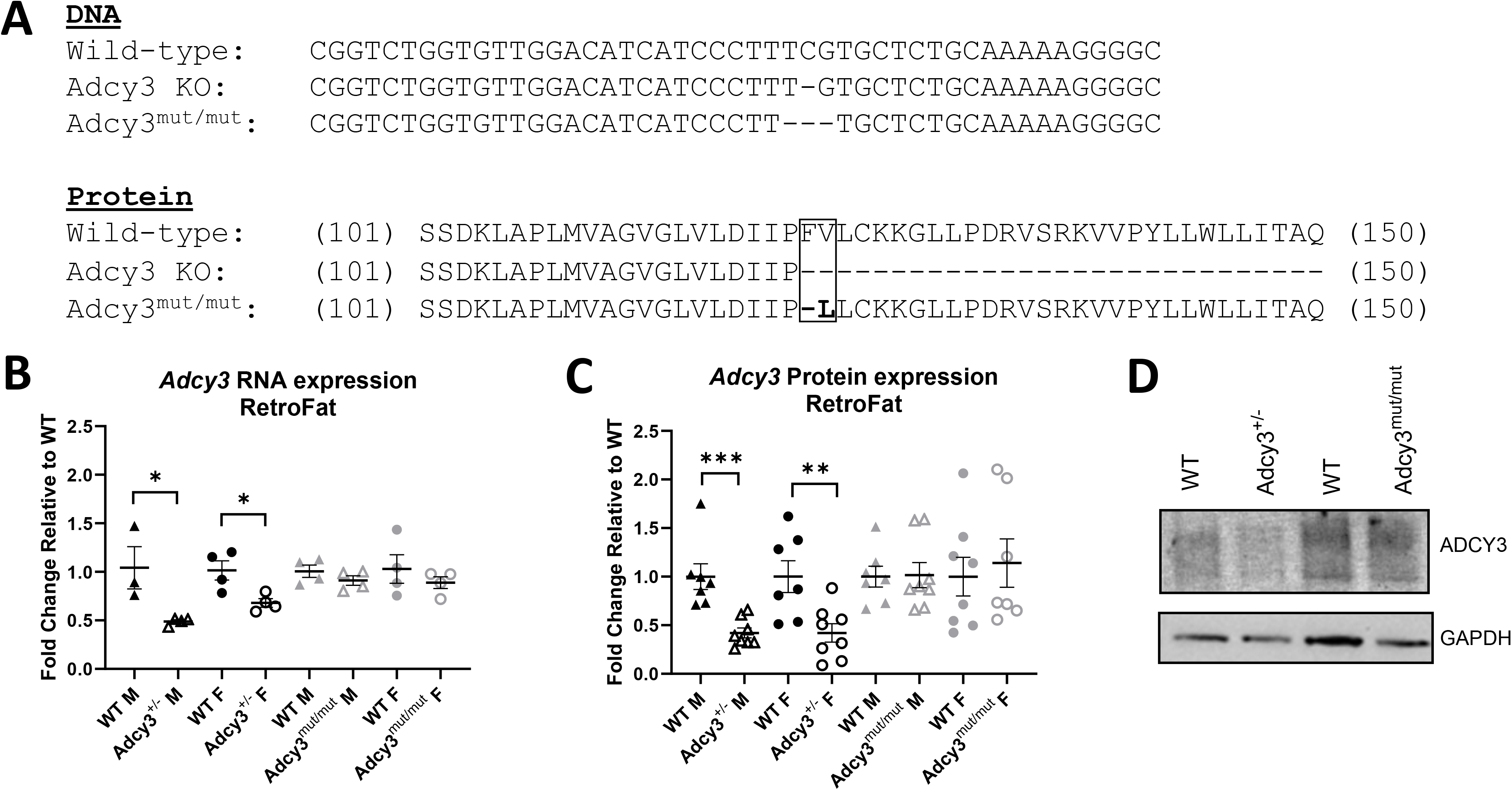
Development of *Adcy3* KO and Adcy3^mut/mut^ rats and confirmation of *Adcy3* expression. **(A)** *Adcy3* knockout (KO) has a single base pair deletion in the transmembrane domain, causing a frameshift mutation and whole-body KO of the gene. Because *Adcy3* KO is lethal, Adcy3^+/-^ rats were used for experiments. Adcy3^mut/mut^ has a 3-base pair deletion in the same transmembrane region, causing a F122delV123L protein-coding mutation. **(B)** Adcy3^+/-^ males (M) and females (F) express less *Adcy3* mRNA than wild type (WT) rats in retroperitoneal white adipose (RetroFat). Adcy3^mut/mut^ does not have altered *Adcy3* mRNA expression in RetroFat. **(C)** M and F Adcy3^+/-^ rats express less ADCY3 in RetroFat than WT rats. Adcy3^mut/mut^ rats do not have altered ADCY3 RetroFat expression. **(D)** Representative Western Blot image comparing ADCY3 expression across genotypes. GAPDH was used as a loading control. Mean ± SEM. T-test, *p<0.05, **p<0.01, ***p<0.001

Because *Adcy3* KO is lethal shortly after birth, heterozygous Adcy3^+/-^ rats from this colony were used as experimental animals. Notably, Adcy3^mut/mut^ does not display this lethality. Adcy3^+/-^ and WT experimental rats were collected separately from Adcy3^mut/mut^ and WT rats, such that comparisons are between WT and Adcy3^+/-^ or Adcy3^mut/mut^ within each strain, not between strains. A total of 10-12 rats per genotype-sex group were initially used for metabolic and behavioral experiments. Due to data loss during early experiments, we added additional cohorts, resulting in 17-20 rats/group for intraperitoneal glucose tolerance test (IPGTT), individual food intake, and TSE PhenoMaster. A separate group of rats (4/group) was used for olfactory habituation-dishabituation experiments. A third group of rats (4-8/group) was used to collect tissues for qPCR and Western Blot. Housing and diet conditions are described in Supplemental Methods. All animal experiments were performed using protocols approved by the Institutional Animal Care and Use Committee at WFUSOM.

### 3.2 Metabolic and Behavioral Study Design

Rats were weaned at 3 weeks of age and pair-housed, and experimental rats all began a high-fat diet (HFD) (60% kcal fat, 20% kcal protein, 20% kcal carbohydrates, ResearchDiet D12492) at 5 weeks of age (**Figure 2A**). Body weight and cage-wide food intake were measured weekly. Behavioral phenotyping began at 6 weeks on diet (WOD), metabolic phenotyping began at 8 WOD, and rats were euthanized after 12 WOD.

**Figure 2.**
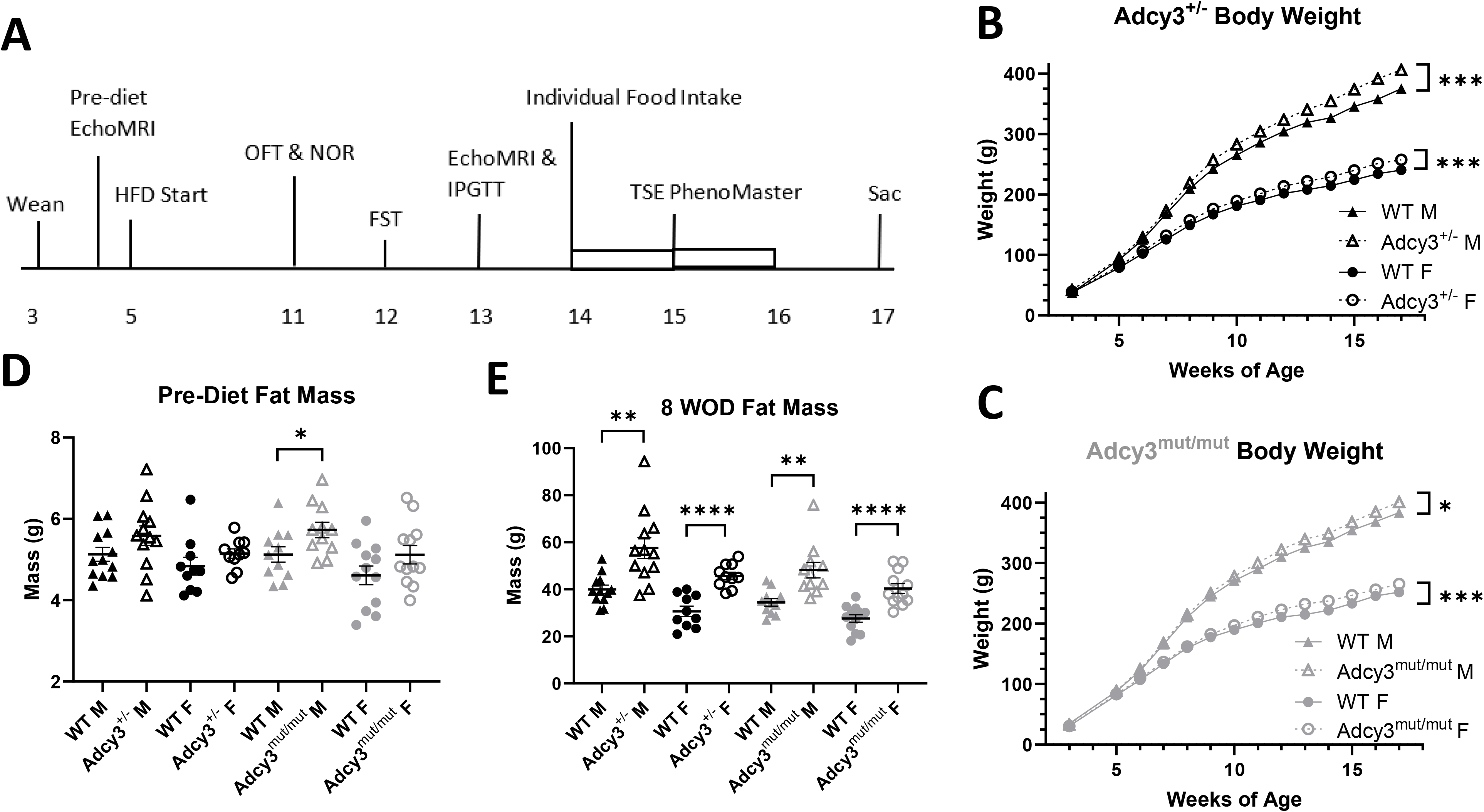
Study design, body weight, and body composition in Adcy3^+/-^ and Adcy3^mut/mut^ rats. **(A)** Timeline of the study shown in weeks of age. Metabolic phenotyping included weekly measurement of body weight and cage-wide food intake, EchoMRI, intraperitoneal glucose tolerance test (IPGTT), individual housing to measure food intake, TSE PhenoMaster chambers, and euthanasia (sac). Behavioral phenotyping included the open field test (OFT), novel object recognition test (NOR), and forced swim test (FST). **(B)** Adcy3^+/-^ males (M) and females (F) gain more weight than wild-type (WT) rats over the course of the study. **(C)** Adcy3^mut/mut^ M and F gain more weight than WT rats over the course of the study. **(D)** There are no differences in fat mass prior to HFD start, except in Adcy3^mut/mut^ M, which have slightly more fat mass than WT M. **(E)** After 8 weeks on diet (WOD), all 4 groups have more fat mass than WT rats. Mean ± SEM. T-test or rmANOVA, *p<0.05, **p<0.01, ***p<0.001, ****p<0.0001

### 3.3 Metabolic Phenotyping

Fat and lean mass were measured using EchoMRI analysis (EchoMRI LLC, Houston, TX) at 4 weeks of age before HFD start, then again after 8 WOD. Rats received an IPGTT after 8 WOD, as described previously^35^ and in Supplemental Methods. After 9 WOD, rats were single-housed for 1 week to measure individual food intake. At 10 WOD, rats spent an additional 5 days single-housed in TSE PhenoMaster indirect calorimetry chambers (TSE Systems, Chesterfield, MO) where oxygen consumption, carbon dioxide production, energy expenditure (EE), locomotor activity, food and water intake, and respiratory exchange ratio (RER) were measured. RER near 0.7 indicates that fat is predominantly being used for energy, while RER near 1.0 indicates carbohydrates are predominant.

### 3.4 Behavioral Phenotyping

All behavior measures were video recorded, and videos were scored by an experimenter blind to the rat’s group. Details of each apparatus and procedure are in Supplemental Methods.

#### 3.4.1 Open Field Test (OFT)

Rats received the OFT after 6 WOD to assess anxiety-like behaviors as previously described.^36^ Measures scored included latency to exit the smaller center circle, time spent in the larger center circle, number of line crossings, number of rearings, number of grooming episodes, and number of fecal boli produced.

#### 3.4.2 Novel Object Recognition Test (NOR)

Rats received the NOR after 6 WOD to assess memory. Briefly, rats were habituated to an object on Day 1 and were presented with this familiar object and a novel object on Day 2. Percentage of time spent with each object was calculated by (time spent with object)/(total time spent with both objects).

#### 3.4.3 Forced Swim Test (FST)

Rats received the FST after 7 WOD to assess passive coping^37^ and/or depression-like behavior.^38^ Rats were placed in a tank of 25 water for 15 minutes on Day 1 and 5 minutes on Day 2. Behavior on Day 2 was scored at 5-second intervals as “mobile” (swimming, climbing, or diving) or “immobile” (floating).

### 3.5 Tissue Harvest and Histology

After 12 WOD, rats were euthanized via live decapitation following a 4-hour fast, and multiple tissues were weighed (Supplemental Methods). Retroperitoneal fat (RetroFat) was collected and stored in 10% formalin for histology as described in Supplemental Methods.

### 3.6 Olfactory Habituation-Dishabituation Test (Anosmia Test)

A separate group of rats received the anosmia test individually in their home cage at 10 weeks of age. The following odorants were presented on a cotton swab (3 times for 2 minutes each): water, almond extract 1:50, opposite-sex rat urine 1:20, peanut butter (1g in 40 ml water). Time spent sniffing the swab at each presentation was scored.

### 3.7 Real-time Quantitative PCR (rt-qPCR) and Western Blot

Total RNA was extracted from RetroFat of rats (4/group) for rt-qPCR analysis, and protein was extracted from RetroFat of rats (7-8/group) for Western Blot. Detailed procedures are provided in Supplemental Methods.

### 3.8 Statistical Analysis

Data were analyzed in RStudio (2023.03.1) and GraphPad Prism (10.0.02). As described in Supplemental Methods, outliers were identified and removed, and data were transformed to fit a normal distribution.

Within each strain (Adcy3^+/-^ or Adcy3^mut/mut^), data were first analyzed using a two-way ANOVA or repeated measures ANOVA (rmANOVA) with genotype and sex as main factors. Because multiple significant sex effects and genotype by sex interactions were observed, data were also stratified by sex and analyzed separately. In the stratified analysis, an unpaired Student’s t-test was used to compare outcome variables for which each rat had a single measurement. A paired Student’s t-test was used to compare time spent with the novel vs. familiar object within each experimental group in the NOR. A one-way rmANOVA was used to compare variables for which repeated measurements were taken (body weight, food intake, and anosmia test). Due to large sex differences in body weight and fat pad weights, we report only the stratified analysis for these phenotypes. TSE PhenoMaster data were analyzed separately for each sex given that different flow rates were used. TSE data were analyzed with the CalR Web Application (version 1.3) using the analysis of covariance (ANCOVA) method as previously described.^39^

## 4 Results

### 4.1 Adcy3^+/-^ rats, but not Adcy3^mut/mut^ rats, have decreased Adcy3 expression

Adcy3^+/-^ males (p=0.0298) and females (p=0.0212) had significantly decreased *Adcy3* mRNA expression in RetroFat, expressing ∼50% *Adcy3* relative to WT (**Figure 1B**). There were no differences in *Adcy3* expression in Adcy3^mut/mut^ compared to WT. Western blots showed similar results for ADCY3 protein expression: Adcy3^+/-^ males (p=0.0009) and females (p=0.0073) had significantly reduced expression of ADCY3 (**Figure 1C, 1D**) while Adcy3^mut/mut^ did not. These results confirm that Adcy3^+/-^ has an expression-altering variant while Adcy3^mut/mut^ has a protein-coding variant that does not alter total ADCY3 expression.

### 4.2 Adcy3^+/-^ and Adcy3^mut/mut^ rats do not show evidence of anosmia

Given that *Adcy3* KO mice have been reported to have anosmia^25^ and that anosmia may affect food intake, we assessed olfaction in Adcy3^+/-^ and Adcy3^mut/mut^. There were no differences in time spent sniffing any of the four odorants in Adcy3^+/-^ or Adcy3^mut/mut^ males or females relative to WT, indicating that olfaction is not impaired (**Figure S2**).

### 4.3 Adcy3^+/-^ and Adcy3^mut/mut^ rats weigh more than WT rats due to increased fat mass

Both Adcy3^+/-^ males (p=0.0001) and females (p=0.0005) gained significantly more weight than WT rats over the course of the study (**Figure 2B**), as did Adcy3^mut/mut^ males (p=0.0280) and females (p=0.0002) (**Figure 2C**). Prior to HFD start, differences in fat mass were minimal (**Figure 2D**). After 8 WOD, Adcy3^+/-^ (p=0.0015; p<0.0001) and Adcy3^mut/mut^ (p=0.0013; p<0.0001) males and females all had significantly more fat mass than WT rats (**Figure 2E**). No changes in lean mass were found at either time point (**Figure S3**).

At euthanasia, Adcy3^+/-^ males and females had significantly more RetroFat (p=0.0003; p=0.0002), gonadal fat (p=0.0002; p<0.0001), and omental fat (OmenFat) (p=0.0019; p=0.0003) than WT rats (**Figure 3A**). Adcy3^mut/mut^ males and females also had significantly more RetroFat (p=0.0060; p<0.0001), gonadal fat (p=0.0080; p<0.0001), and omental fat (OmenFat) (p=0.0059; p=0.0026) than WT rats (**Figure 3B**). Adcy3^+/-^ females had slightly heavier pancreases than WT females (p=0.0246) (**Figure S3**). There were no differences in the weights of any other tissues (**Figure S3**). Adcy3^+/-^ (p=0.0159; p<0.0001) and Adcy3^mut/mut^ (p<0.0001, p<0.0001) males and females all had significantly larger adipocytes in RetroFat than WT rats (**Figure 3C-E**).

**Figure 3.**
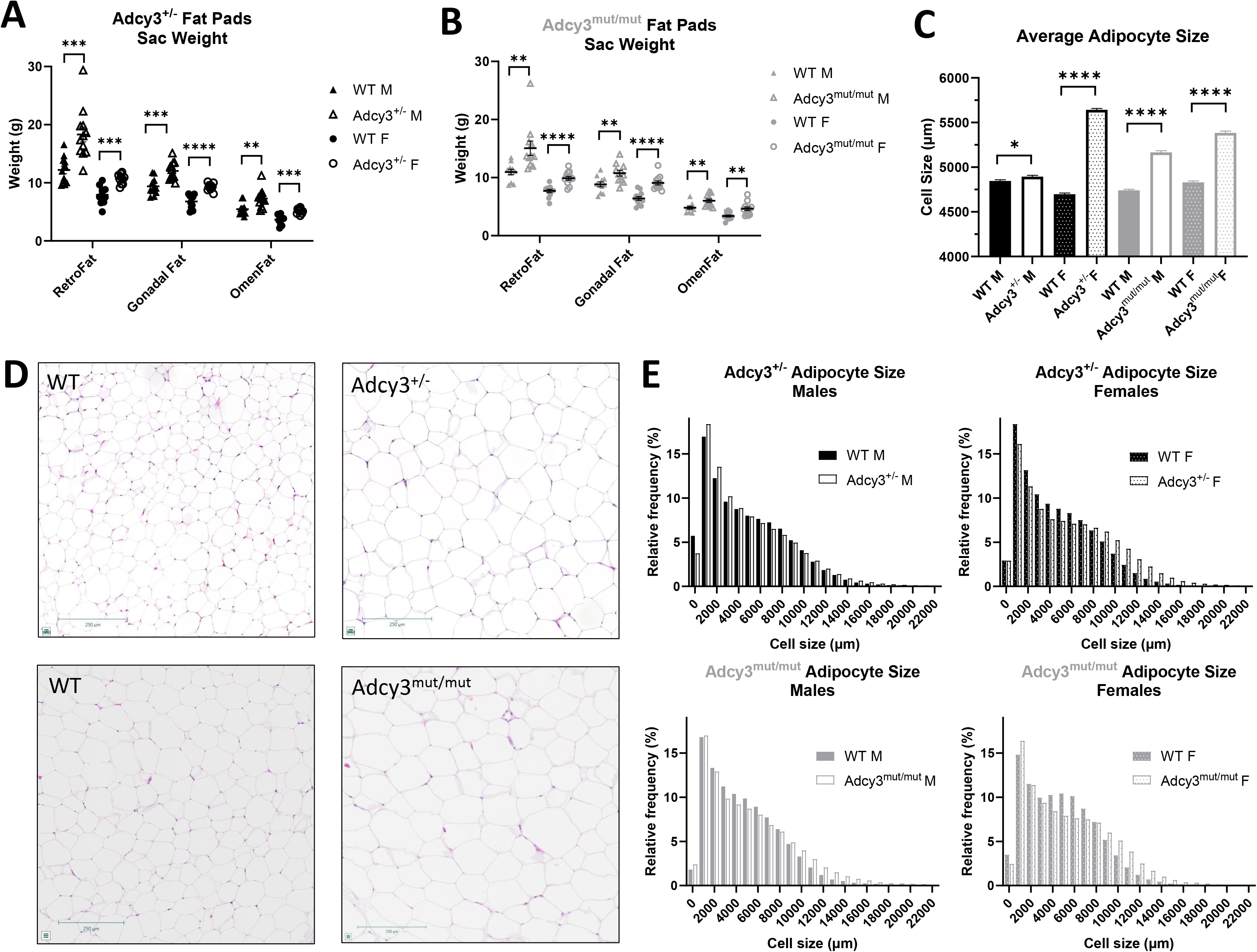
Adcy3^+/-^ and Adcy3^mut/mut^ fat pads and adipocytes at sac. **(A)** Adcy3^+/-^ males (M) and females (F) have larger retroperitoneal (RetroFat), gonadal, and omental (OmenFat) fat pads at sac than wild-type (WT). **(B)** Adcy3^mut/mut^ M and F also have larger RetroFat, gonadal fat, and OmenFat pads at sac than WT. **(C)** Adcy3^+/-^ and Adcy3^mut/mut^ M and F all have significantly larger adipocytes than WT. **(D)** Representative images of hematoxylin and eosin-stained RetroFat from female rats. Scale bars = 250μm. **(E)** Relative frequency distributions of adipocyte size for each group. Mean ± SEM. T-test, *p<0.05, **p<0.01, ***p<0.001, ****p<0.0001

### 4.4 Adiposity is driven by decreased energy expenditure in females, but by increased food intake in males

In the TSE PhenoMaster chambers, relative to WT females, Adcy3^+/-^ females had significantly decreased average daily EE (p=0.0068) and cumulative EE (p=0.0265) (**Figure 4B**). There were no differences in EE in Adcy3^+/-^ males (**Figure 4A**). Similarly, Adcy3^mut/mut^ females had significantly decreased average daily EE (p=0.0264) and cumulative EE (p=0.0324) (**Figure 4D**) while there were no differences in EE in Adcy3^mut/mut^ males (**Figure 4C**). Adcy3^+/-^ males (p=0.0100) (**Figure 4A**), but not females (**Figure 4B**), had a significantly increased RER relative to WT rats. There were no differences in RER in Adcy3^mut/mut^ (**Figure 4C, 4D**). There were no differences in water consumption or locomotor activity between groups (**Figure S4**).

**Figure 4.**
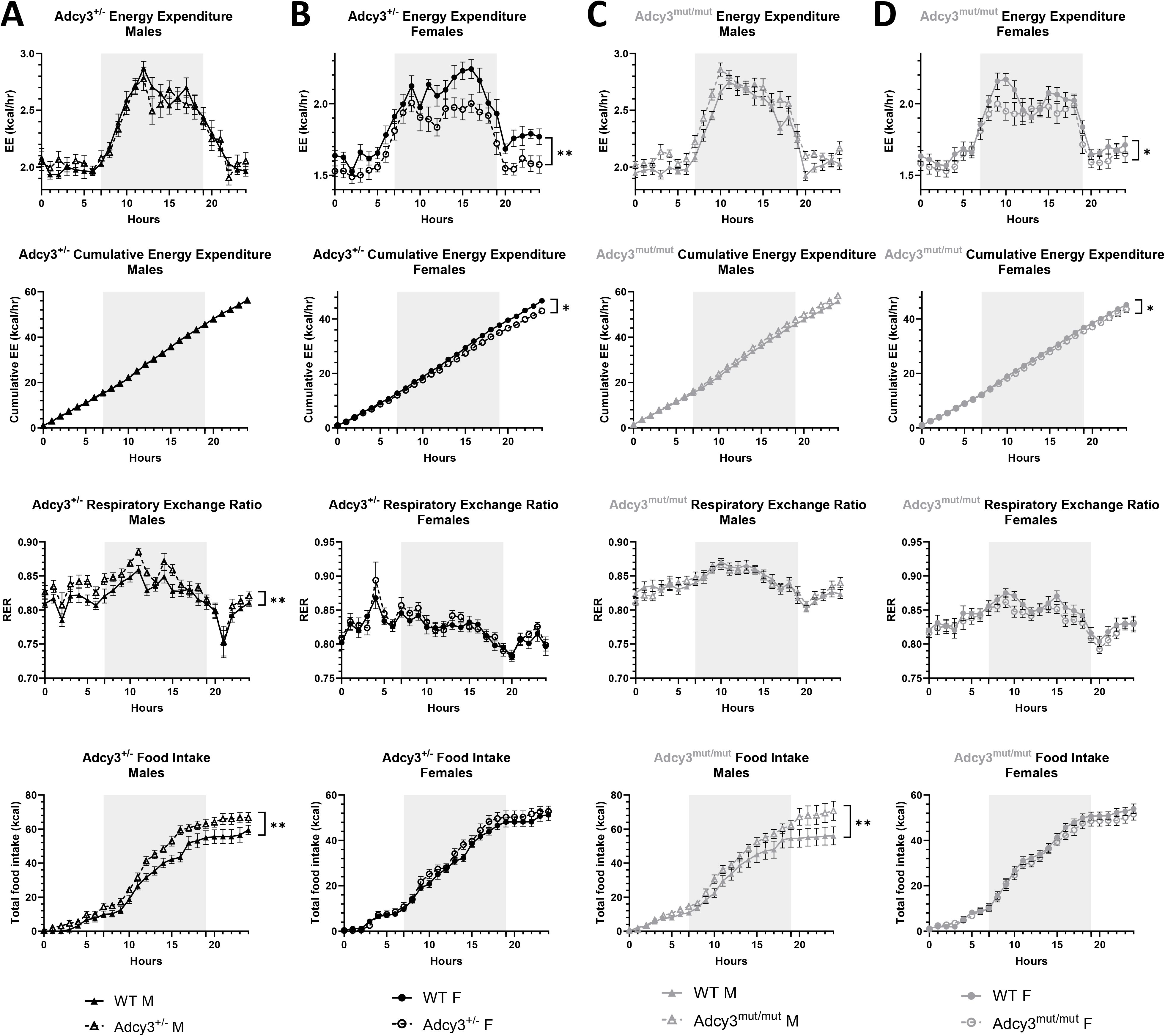
Sex differences in the cause of adiposity in Adcy3^+/-^ and Adcy3^mut/mut^ as measured in TSE PhenoMaster chambers. Plots of 24 hours of average hourly energy expenditure (EE), cumulative EE, respiratory exchange ratio (RER), and cumulative food intake in **(A)** Adcy3^+/-^ males (M), **(B)** Adcy3^+/-^ females (F), **(C)** Adcy3^mut/mut^ M, and **(D)** Adcy3^mut/mut^ F compared to wild-type (WT). Adcy3^+/-^ and Adcy3^mut/mut^ M have increased food intake, while Adcy3^+/-^ and Adcy3^mut/mut^ F have decreased EE. Only Adcy3^+/-^ M had increased RER. Gray shading indicates dark cycle. Mean ± SEM. ANCOVA, *p<0.05, **p<0.01

Adcy3^+/-^ males consumed significantly more food than WT males in the TSE chambers (p=0.0075) (**Figure 4A**) while there were no differences in food intake in Adcy3^+/-^ females (**Figure 4B**). Adcy3^mut/mut^ males also consumed significantly more food than WT males (p=0.0079) (**Figure 4C**) while there were no differences in Adcy3^mut/mut^ females (**Figure 4D**). We also measured food intake in the rats’ normal caging in two ways: cage-wide intake on a weekly basis, and individual intake over one week. For cage-wide and individual food intake, despite effects of sex, there were no genotype effects and no sex by genotype interaction in Adcy3^+/-^ (**Figure 5A-C**). In Adcy3^mut/mut^ rats, however, for cage-wide intake, there was significant three-way interaction between sex, genotype, and time (p=0.0430), where Adcy3^mut/mut^ males consumed significantly more food than WT males over time (p=0.0210) (**Figure 5D**) with no differences in Adcy3^mut/mut^ females (**Figure 5E**), supporting the TSE chamber findings. For individual intake, there were significant effects of sex (p<0.0001) and genotype (p=0.0336), where Adcy3^mut/mut^ males consumed significantly more food than WT males (p=0.0425) with no differences in females (**Figure 5F**).

**Figure 5.**
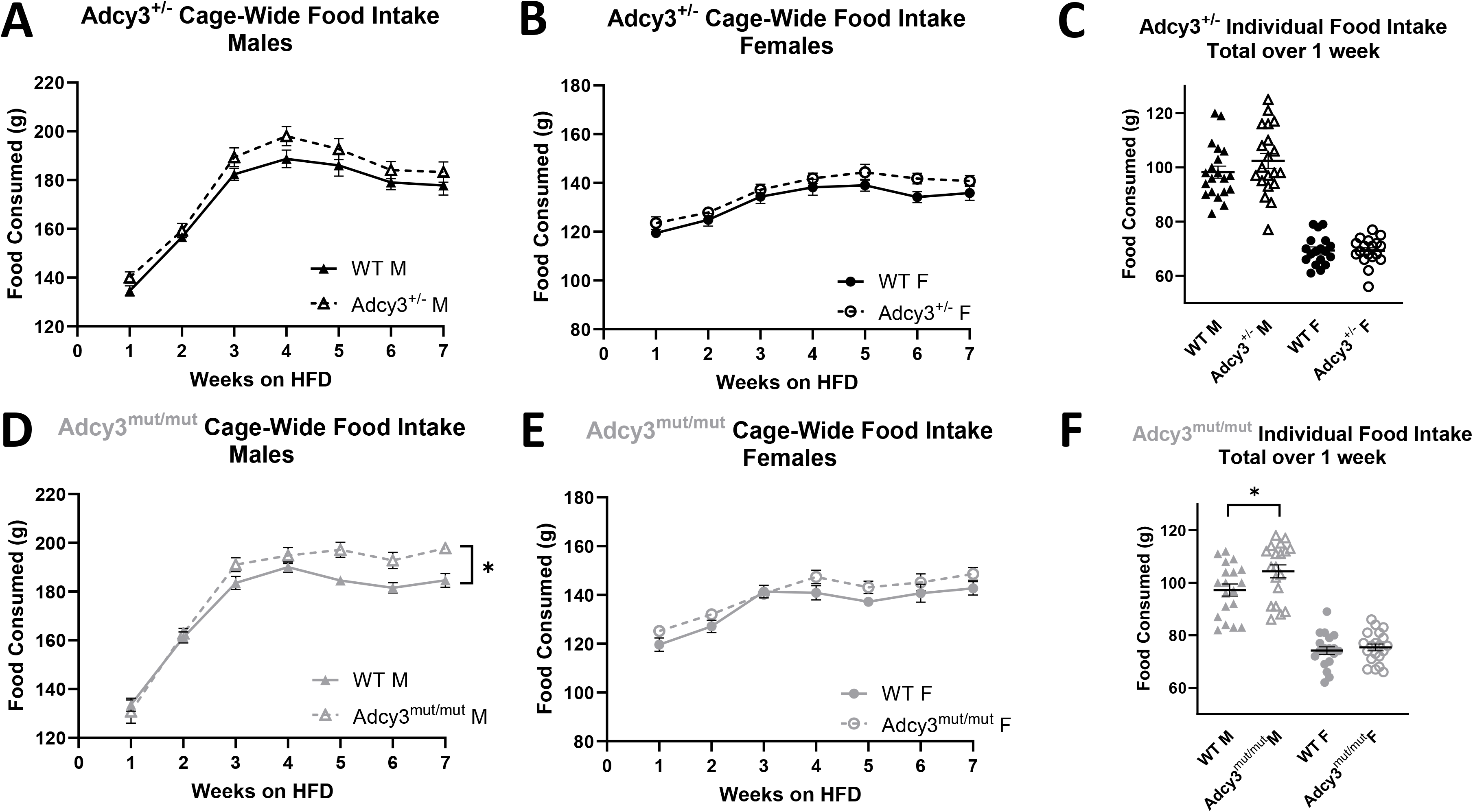
Cage-wide and individual food intake in Adcy3^+/-^ and Adcy3^mut/mut^. **(A)** No differences in cage-wide weekly food intake in Adcy3^+/-^ males (M) **(B)** or Adcy3^+/-^ females (F). **(C)** No differences in individual food intake over one week in Adcy3^+/-^ M or F. **(D)** Adcy3^mut/mut^ M consume more food than wild-type (WT) M. **(E)** No differences in cage-wide weekly food intake in Adcy3^mut/mut^ F. **(F)** Adcy3^mut/mut^ M, but not F, consume more food than WT when measured individually over one week. HFD: high-fat diet. Mean ± SEM. T-test or rmANOVA, *p<0.05

### 4.5 Adcy3^+/-^ and Adcy3^mut/mut^ males are more insulin resistant than WT males

In Adcy3^+/-^ rats, there were significant sex and genotype effects for fasting glucose (p=0.0076; p=0.0311), fasting insulin (p=0.0387; p=0.0179), and HOMA-IR (p=0.0084; p=0.0107). There were also significant two-way interactions for fasting insulin (p=0.0402) and HOMA-IR (p=0.0308). Adcy3^+/-^ males had significantly higher fasting glucose (p=0.0297) (**Figure 6A**), fasting insulin (p=0.0031) (**Figure 6B**), and HOMA-IR (p=0.0028) (**Figure 6C**) than WT males, with no differences in these measures between Adcy3^+/-^ and WT females. In Adcy3^mut/mut^ rats, there were significant genotype effects for fasting insulin (p=0.0121) and HOMA-IR (p=0.0171). When separated by sex, Adcy3^mut/mut^ males had significantly higher fasting insulin (p=0.0173) (**Figure 6B**) and HOMA-IR (p=0.0276) (**Figure 6C**) than WT males, with no significant differences between Adcy3^mut/mut^ and WT females. In both Adcy3^+/-^ and Adcy3^mut/mut^, there were no differences in glucose AUC between genotypes (**Figure S2**).

**Figure 6.**
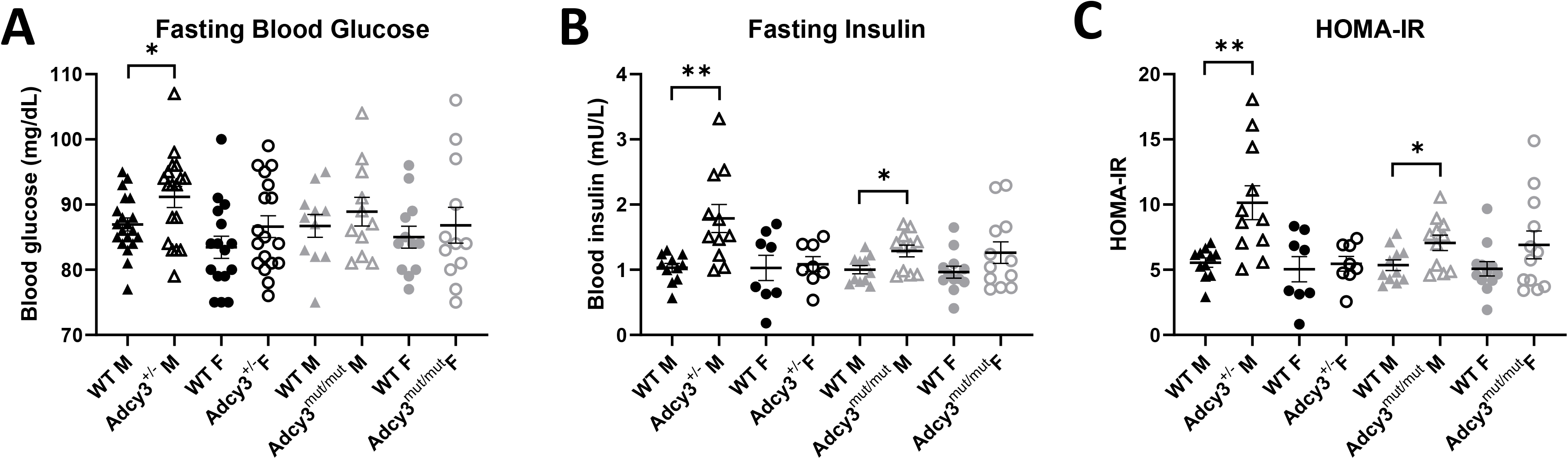
Measures of glucose tolerance and insulin resistance in Adcy3^+/-^ and Adcy3^mut/mut^. **(A)** Only Adcy3^+/-^ males (M) have increased blood glucose compared to wild-type (WT) after an overnight fast. **(B)** Adcy3^+/-^ and Adcy3^mut/mut^ M have increased fasting insulin after an overnight fast, with no differences in females (F). **(C)** Adcy3^+/-^ and Adcy3^mut/mut^ M have increased insulin resistance compared to WT as measured by HOMA-IR. There were no differences in HOMA-IR in F. Mean ± SEM. T-test, *p<0.05, **p<0.01

### 4.6 Adcy3^mut/mut^ females show increased anxiety-like behavior in the OFT

In Adcy3^+/-^ rats there was a significant effect of sex for line crossings (p=0.0002) and center time (p=0.0001), where females had increased line crossings and decreased center time relative to males. There were no genotype or interaction effects for any OFT variable in the 2-way ANOVA or when stratified by sex (**Table S2**).

In contrast, for center time in Adcy3^mut/mut^, there were significant effects of both sex (p=0.0117) and genotype (p=0.0427) although there was no interaction effect. When separated by sex, however, Adcy3^mut/mut^ females spent significantly less time in the center of the arena (p=0.0221) than WT females while Adcy3^mut/mut^ males showed no differences in center time (**Figure 7A**). For rearing episodes, there was a significant effect of sex (p=0.0002) and a significant two-way interaction between sex and genotype (p=0.0025), where Adcy3^mut/mut^ females spent significantly less time rearing (p=0.0220) than WT females, but Adcy3^mut/mut^ males spent significantly *more* time rearing than WT males (p=0.0495) (**Figure 7B**). For grooming episodes, there was a significant two-way interaction effect between sex and genotype (p=0.0320) where Adcy3^mut/mut^ males tended to spend more time grooming (p=0.0830) than WT males, with no differences in grooming between Adcy3^mut/mut^ and WT females (**Figure 7C**). There was a significant effect of sex on latency to leave the center (p=0.0072), fecal boli (p=0.0097), and line crossings (p<0.0001) without any genotype or interaction effects. When separated by sex, there were no significant differences between genotypes in these three variables (**Figure 7D, Table S2**). Collectively, these data are consistent with an increase in anxiety-like behaviors in Adcy3^mut/mut^ females while the pattern of behavior in males is less clear.

**Figure 7.**
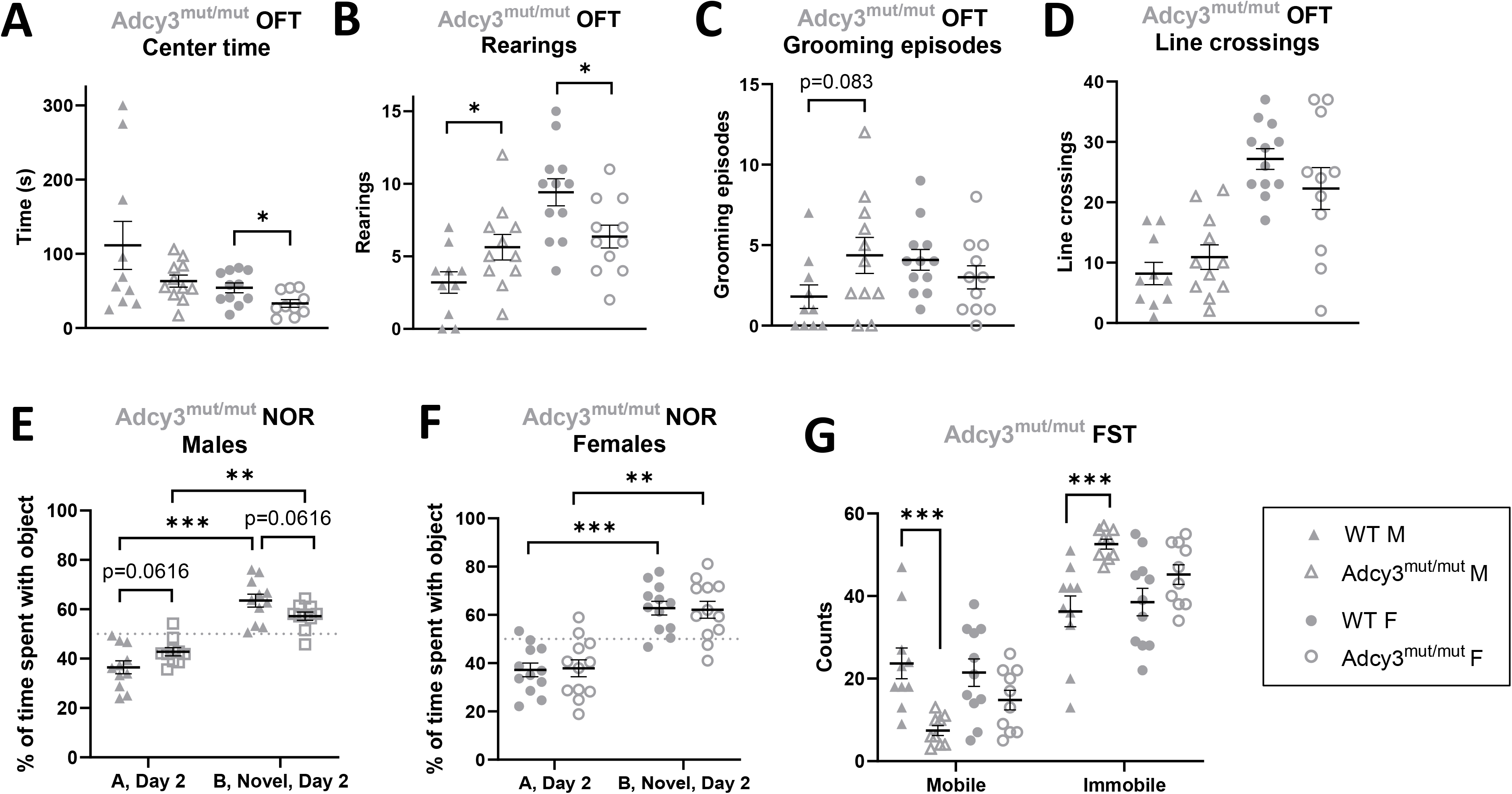
Adcy3^mut/mut^ behaviors in the open field test (OFT), novel object recognition test (NOR), and forced swim test (FST). **(A)** Adcy3^mut/mut^ females (F), but not males (M), spend significantly less time in the center than wild-type (WT). **(B)** Adcy3^mut/mut^ M rear significantly more and Adcy3^mut/mut^ F rear significantly less than WT. **(C)** Adcy3^mut/mut^ M tend to groom more than WT M. **(D)** There were no differences in line crossings in Adcy3^mut/mut^ M or F. **(E)** “A”: familiar object, “B”: novel object. Adcy3^mut/mut^ and WT M both spend significantly more time with the novel object than the familiar object, but Adcy3^mut/mut^ M tend to spend less time with the novel object compared to WT M, potentially indicating impaired memory. **(F)** Although Adcy3^mut/mut^ F also spend more time with the novel object than with the familiar object, there were no differences in NOR behavior compared to WT. **(G)** Adcy3^mut/mut^ M spend more time immobile in the FST than WT M. There were no differences in behavior in Adcy3^mut/mut^ F. Mean ± SEM. T-test, *p<0.05, **p<0.01, ***p<0.001

### 4.7 Adcy3^mut/mut^ males show decreased memory in the NOR

No significant sex, genotype, or interaction effects were observed for any group or variable in the NOR when analyzed with a 2-way ANOVA. As expected, on Day 2, both WT (p=0.0025; p<0.0001) and Adcy3^+/-^ (p<0.0001; p<0.0001) males and females spent significantly more time with the novel object vs. the familiar object (**Table S2**). There were no differences in time spent with the novel object between genotypes (**Table S2**). Similarly, on Day 2, WT (p=0.0004; p=0.0009) and Adcy3^mut/mut^ (p=0.0019; p=0.0054) males and females all spent significantly more time with the novel object vs. the familiar object (**Figure 7E-F**). However, when stratified by sex, Adcy3^mut/mut^ males tended to spend a smaller percentage of the time with the novel object relative to WT males (p=0.0616), which may indicate decreased memory for the familiar object (**Figure 7E**). There were no differences in the percentage of time spent with the novel object between WT and Adcy3^mut/mut^ females (**Figure 7F**). We note that on Day 1, when rats received two identical objects in their cage, WT males from the Adcy3^mut/mut^ strain spent more time with the object at the back of the cage while all other groups spent approximately 50% of the time at each side of the cage (**Figure S5, Table S2**).

### 4.8 Adcy3^mut/mut^ males show increased passive coping in the FST

In Adcy3^+/-^, there were significant effects of sex on immobile and mobile behavior counts (p=0.0027), where females were more mobile compared to males. However, there were no genotype effects in the stratified analysis (**Table S2**).

In Adcy3^mut/mut^, there were significant effects of genotype on immobile and mobile behavior counts (p=0.0004) but no sex or interaction effects. When separated by sex, however, Adcy3^mut/mut^ males spent significantly more time immobile and significantly less time mobile than WT males (p=0.0010), consistent with an increase in passive coping (**Figure 7G**). There were no differences in FST behavior between Adcy3^mut/mut^ and WT females (**Figure 7G**).

## 5 Discussion

We demonstrate that Adcy3^mut/mut^ rats have increased body weight due to increased fat mass, as well as larger adipocytes, relative to WT rats. This phenotype aligns with what we see in Adcy3^+/-^ rats, underscoring the importance of the TM domain for ADCY3 function in adiposity. We further demonstrate that the underlying cause of adiposity differs between male and female rats. Adiposity increased in males due to increased food intake and in females due to decreased energy expenditure. Finally, we show that Adcy3^mut/mut^ males, but not females, show increased passive coping in the FST and possibly decreased memory in the NOR, two phenotypes associated with despair-like behavior. Adcy3^mut/mut^ females, on the other hand, do not show differences in FST or NOR but do show increased anxiety-like behaviors in the OFT. This is the first study to demonstrate the impact of a protein-coding mutation in *Adcy3* on adiposity and behavior in a rodent model, as well as the first to report the importance of sex differences for both phenotypes.

The fact that we observed similar body weight and adiposity phenotypes in Adcy3^mut/mut^ and Adcy3^+/-^ rats, consistent with previous findings in Adcy3 KO mice,^22–24^ demonstrates that the ADCY3 TM domain where Adcy3^mut/mut^ is located plays a key role in protein function. Although the TM domain of ADCY3 has previously been identified as important in membrane localization^40^ and catalytic efficiency,^41^ and obesity-associated variants in this domain have been identified in humans,^13^ this is the first study to directly investigate the *in vivo* effects of a TM domain mutation.

The underlying cause of adiposity in both Adcy3^+/-^ and Adcy3^mut/mut^ rats differed by sex. Female Adcy3^+/-^ and Adcy3^mut/mut^ rats expended less energy than WT rats without any changes in food intake. In contrast, male Adcy3^+/-^ and Adcy3^mut/mut^ rats had no changes in EE but had increased food intake relative to WT in the TSE chambers. Interestingly, outside of the TSE chambers, we observed increased cage-wide and individual food intake in Adcy3^mut/mut^ males but not in Adcy3^+/-^ males, suggesting that either two altered copies of *Adcy3* or an additional stressor (such as the stress of the novel TSE environment) is necessary to observe effects on food intake. Our findings are consistent with previous studies showing *Adcy3* KO male mice consume more food than WT,^23, 24^ while the few studies that have investigated females reported no changes in food intake.^22, 24^ EE had not been previously assessed in any *Adcy3* KO model, but one study examining a gain-of-function mutation in *Adcy3* reported that EE played a role in protecting against diet-induced obesity, although this study was only done in male mice.^27^

The difference in what is causing adiposity in males vs. females might be explained by the influence of sex hormones in the hypothalamus and in adipose tissue. *Adcy3* is expressed in both of these tissues,^42^ and estrogen in particular has been shown to act in the hypothalamus to influence thermogenesis, adipose distribution, and brain-adipose crosstalk.^43^ It has also been shown that obesity-associated genetic variants in humans can have differing, even entirely opposite, effects in men vs. women.^44^ Our future studies will examine the molecular mechanisms of these observed sex differences in more detail, but the current study emphasizes the importance of assessing both sexes in obesity research as the specific phenotype (as well as potential therapies) may be remarkably different.

We also found that Adcy3^+/-^ and Adcy3^mut/mut^ males, but not females, had increased insulin resistance compared to WT rats. These results were unexpected given that both sexes had increased body weight and fat mass. The effect of sex was found in Adcy3^mut/mut^ rats only in the stratified analysis, so it will be important to repeat these studies with a larger number of animals to validate this finding. The fact that we saw an effect only in males in both strains, however, suggests the sex effect is real. Sex differences in insulin resistance in response to a HFD have been documented in the literature,^45^ although the underlying cause of these differences remains an open question.

We also observed altered behavior in both Adcy3^mut/mut^ males and females, and we again observed sex differences. The strongest phenotype we observed was in the FST, where Adcy3^mut/mut^ males showed significantly increased immobility relative to WT, indicating increased passive coping and/or despair-like behavior.^38^ This finding is consistent with previous findings in male *Adcy3* KO mice.^25^ Adcy3^mut/mut^ males also showed a trend towards impaired memory in the NOR, as previously shown for *Adcy3* KO mice.^25^ Both impaired memory and increased passive coping are consistent with a “depression-like” phenotype.^25^ Adcy3^mut/mut^ male OFT behavior was more difficult to interpret as they spent *more* time rearing, but also more time grooming, with no differences in center time, differing from previous findings in *Adcy3* KO mice^25, 28^ and potentially indicating anxious hyperactivity. The behavioral phenotypes we saw in females differed: there were no significant differences in the FST or NOR relative to WT, but in the OFT, Adcy3^mut/mut^ females showed a classic anxiety-like phenotype: spending less time in the center and less time rearing. Collectively, the behavior test results indicate that a TM-domain mutation in *Adcy3* impacts emotional behaviors differently dependent on sex, although more robust testing is needed to fully understand these differences. While it has been suggested that reduced neuronal activity may cause behavior changes in *Adcy3* KO mice^25^, very little is known about the actual underlying mechanisms or brain regions, and more research is needed.

In behavior tests, we unexpectedly did not observe any altered behaviors in Adcy3^+/-^ rats. Previous studies that have examined behavior in *Adcy3* KO models have used models where both copies of the gene are damaged.^25^ Therefore, like the food intake results, these results suggest that one functioning copy of *Adcy3* is sufficient to protect against behavioral effects in rodents.

In conclusion, this work demonstrates four central points. First, protein coding mutations in *Adcy3*, especially in the TM domain, can critically affect protein function to produce similar phenotypes to *Adcy3* KO models. Second, the same genetic changes in *Adcy3* can underlie both adiposity and behavior phenotypes, supporting the idea that *Adcy3* may be a genetic link between obesity and major depressive disorder (MDD)^29^. Third, the underlying cause for adiposity in both mutant and KO models differ by sex. Finally, an *Adcy3* mutation affected emotion-like behaviors differently in male and female rats. These data underscore the importance of continuing to assess both sexes, in both human and rodent studies, in obesity and MDD research. Future studies on Adcy3^mut/mut^ will more precisely investigate the molecular mechanisms underlying these phenotypes.

## Acknowledgments

The authors thank the WFUSOM Comparative Pathology Core, WFUSOM Virtual Microscopy Core, Medical College of Wisconsin Rodent Model Resource, and the Rat Models and Genotyping Service Center for their assistance.

