## Supplemental File for "Protein-coding mutation in *Adcy3* increases adiposity and alters emotional behaviors sex-dependently in rats"

**Contact Info**

Dr. Leah Solberg Woods

575 Patterson Ave, Winston-Salem, NC, 27101;

### Supplemental Methods

*S1.1 Animals and Diet*

*Adcy3* knockout (KO) and Adcy3$^{mut/mut}$ founder animals were genotyped by fragment analysis genotyping assay and confirmed by Sanger sequencing. Founders were then backcrossed to the WKY parental strain, and heterozygous breeding pairs were used to establish *Adcy3* KO and Adcy3$^{mut/mut}$ breeding colonies at the Wake Forest University School of Medicine (WFUSOM). All experimental rats from the WFUSOM breeding colonies were genotyped at the

Medical College of Wisconsin using fluorescent fragment analysis with an ABI 3730 capillary sequencer and Genemapper software (**Table S1**).

All rats were housed at standard temperature and humidity conditions on a 12-hour light/dark cycle with *ad libitum* access to food and water. Food for all breeders, olfactory habituation-dishabituation rats, and rt-qPCR/western blot tissue rats consisted of standard rodent chow (Lab Diet, Prolab RMH 3000, #5P00). Food for metabolic/behavioral experimental rats up to 5 weeks of age also consisted of standard rodent chow. Because some studies show that *Adcy3* KO females only develop increased adiposity when on a high-fat diet[1], at 5 weeks of age, all metabolic/behavioral experimental rats began a high-fat diet (HFD) (60% kcal fat, 20% kcal protein, 20% kcal carbohydrates, ResearchDiet D12492).

#### *S1.2 EchoMRI*

Rats were weighed prior to EchoMRI analysis. To measure fat mass and lean mass, rats were then scanned for 2 minutes in triplicate.

#### *S1.3 Intraperitoneal Glucose Tolerance Test (IPGTT)*

Rats were fasted overnight for 16 hours, then they were given an IP injection of glucose at 1 mg/kg body weight. Blood glucose was measured (Contour Next EZ) and serum was collected from a tail snip at 0, 15, 30, 60, 90, and 120 minutes after injection. Serum insulin was measured with an ultrasensitive ELISA kit (ALPCO, #80-INSRTU-E10). Glucose sensitivity was calculated by measuring area-under-the-curve (AUC) where:

$$AUC = \sum \frac{(S_t + S_{t+1}) \times \Delta min}{2}$$

Homeostatic model assessment for insulin resistance (HOMA-IR) was also calculated to approximate insulin resistance where:

$$HOMA-IR = \frac{fasting\ glucose\left(\frac{mg}{dL}\right) \times fasting\ insulin\ (\frac{mU}{L})}{405}$$

*S1.4 Behavioral Phenotyping*

Any shared items (arena, objects, etc.) were disinfected with 70% ethanol between rats. Rats were given 30 minutes to acclimate to the testing room before each behavioral test.

S1.4.1 Open Field Test (OFT)

OFT was conducted in an open field box (60 x 66 x 40 cm) with black sides and floor under standard room lighting. Concentric circles were drawn on the floor of the box for use in scoring as previously described.[2] Each rat was placed in the center of the box, and behavior was recorded for 5 minutes.

S1.4.2 Novel Object Recognition Test (NOR)

NOR was conducted in the rat's home cage surrounded by black poster board to prevent the rat from looking or climbing out of the cage. During habituation (Day 1), each rat was acclimated to the cage setup for 10 minutes. Then, during familiarization (also Day 1), the rat was given two identical objects (either soda cans or block towers) placed on opposite ends of the home cage and had 10 minutes to freely investigate. 24 hours later (Day 2), during testing, the rat was given one familiar object from the day before and the other, novel object and was given 5 minutes to freely investigate. "Investigating" was defined as the rat directing their nose and attention towards the object while simultaneously having their nose within 1 cm of the object.

Counterbalancing was used for the identity of the novel object and for the cage side on which the novel object appeared (e.g. half the time the novel object was at the front of the cage and half the time it was at the back of the cage).

S1.4.3 Forced Swim Test (FST)

Each rat was placed in a cylindrical Plexiglass tank (30 cm diameter x 50 cm tall) filled with 21 liters of 25℃ water.

*S1.5 Tissue Harvest and Histology*

The following tissues were weighed and snap frozen in liquid nitrogen: brain, pituitary, liver, retroperitoneal fat pads (RetroFat), gonadal fat, gonads, adrenal glands, spleen, brown adipose, subcutaneous adipose, tail. Kidneys, heart, pancreas, and omental fat (OmenFat) were weighed.

RetroFat histology samples were fixed in formalin for 1 week before moving to 70% ethanol for storage. Samples (N=6 per group) were sliced, mounted, and stained with hematoxylin and eosin by the Comparative Pathology Core. Slides were imaged by the Virtual Microscopy Core, then analyzed in Visiopharm (Hoersholm, Denmark). The areas of at least 3000 adipocytes were measured per slide (1 slide per rat), and adipocyte size data was then pooled within each group. Only adipocytes 20$\mu m^2$ - 22,000 $\mu m^2$ were included in the analysis.

*S1.6 Real-time Quantitative PCR (rt-qPCR)*

RNA extraction for rt-qPCR analysis was performed using the RNeasy Lipid Tissue Mini Kit (Qiagen, #74804). mRNA was quantified and assessed for purity (260/280 ratio) on the

Nanodrop Lite Spectrophotometer (Thermo Scientific). mRNA was converted to cDNA using the High-Capacity cDNA Reverse Transcription Kit (Applied Biosystems, 4368814). Rt-qPCR was performed with PowerUp SYBR Green Master Mix (Applied Biosystems, A25741). *β-actin* (Integrated DNA Technologies, **Table S1**) was used as a reference gene. Fold change was calculated using the $2^{-\Delta\Delta Ct}$ method[3] where ΔCt is the difference between the cycle threshold (Ct) of *Adcy3* (Integrated DNA Technologies**, Table S1**) and *β-actin*, and ΔΔCt is the difference between sample ΔCt and the average ΔCt of all WT rats.

*S1.7 Western Blot*

Protein extraction from RetroFat was performed with Pierce IP lysis buffer (Thermo Scientific, #87787), cOmplete Protease Inhibitor Cocktail (Roche, #04693159001), and Bead Beater Tissue Homogenizer at 2200rpm with 1.0mm zirconia/silica beads (BioSpec, #11079110Z). Protein concentrations were measured by BCA assay (Thermo Scientific, #23225). Total proteins (20µg) were separated on 4-20% Criterion TGX Gels (BioRad, #5671094) and transferred to nitrocellulose membranes. Membranes were incubated overnight at 4°C with primary antibodies to ADCY3 (EnCor, RPCA-ACIII, 1:2000) or GAPDH (Santa Cruz, sc-32233, 1:1000). Secondary antibodies were applied for 1 hour 15 minutes at room temperature. Bands were detected by SuperSignal West Pico PLUS chemiluminescent substrate (Thermo Scientific, #34580) and quantified using in ImageJ.

*S1.8 Statistical Analysis*

Outliers were identified by Grubb's Test, or if multiple outliers were suspected, by the ROUT method in Prism with Q=1%. Data were assessed for normality with the Shapiro-Wilk test and were transformed if not normal. If common transformations did not successfully normalize the data, the Mann-Whitney U test was used in place of a Student's t-test for analysis. For rmANOVAs, sphericity was assessed with Mauchly's Test for Sphericity, and if $p<0.05$, Greenhouse-Geisser correction was applied.

### Supplemental Figures

**Figure S1. Full western blot gels corresponding to representative image in Figure 1D. Red boxes indicate sections used for Figure 1D.**

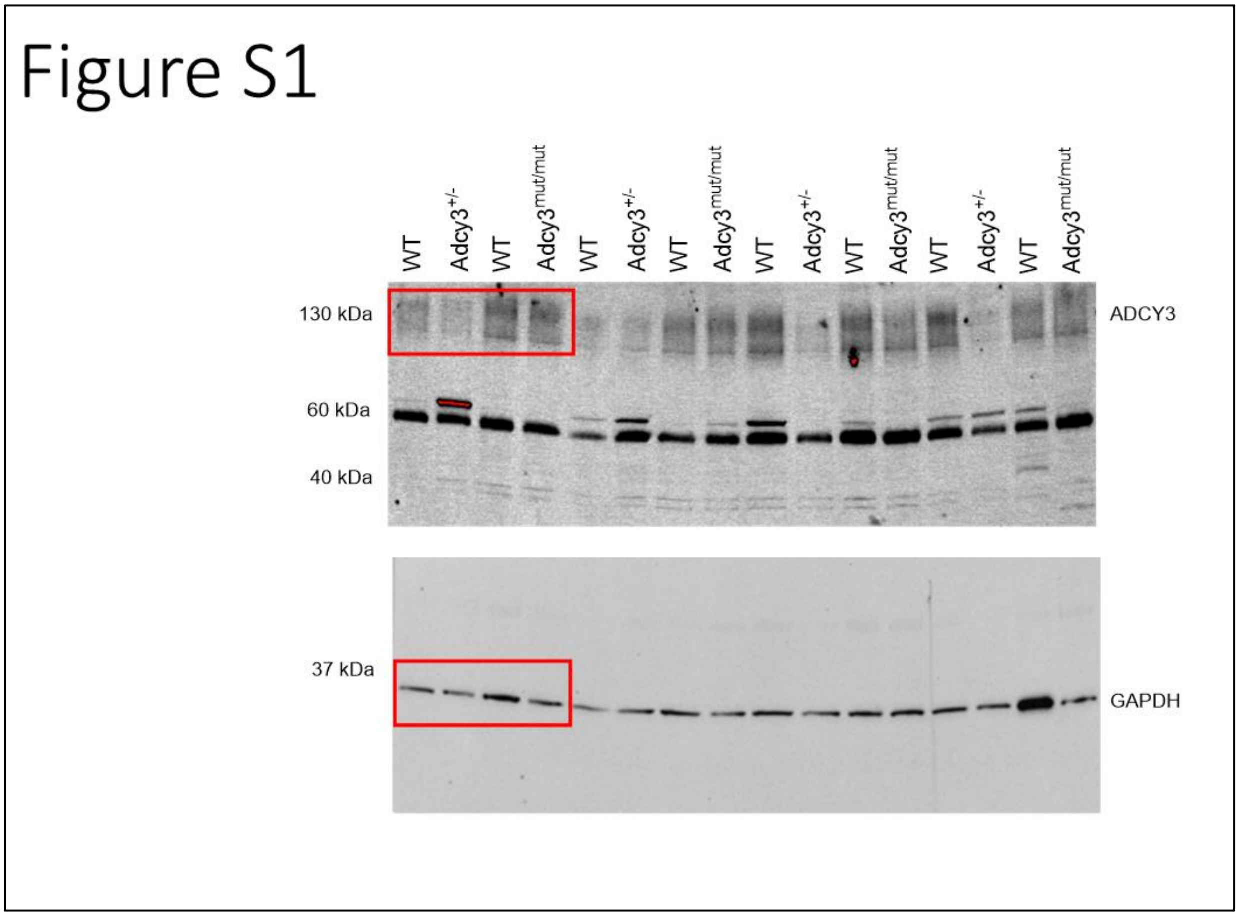

**Figure S2. Olfactory habituation-dishabituation test (anosmia test) in Adcy3$^{+/-}$ and Adcy3$^{mut/mut}$.** No differences in behavior in the anosmia test in any group. W: water, A: almond extract (1:50 in water), U: opposite-sex rat urine (1:20 in water), P: peanut butter (1g in 40 ml water). Mean ± SEM. rmANOVA.

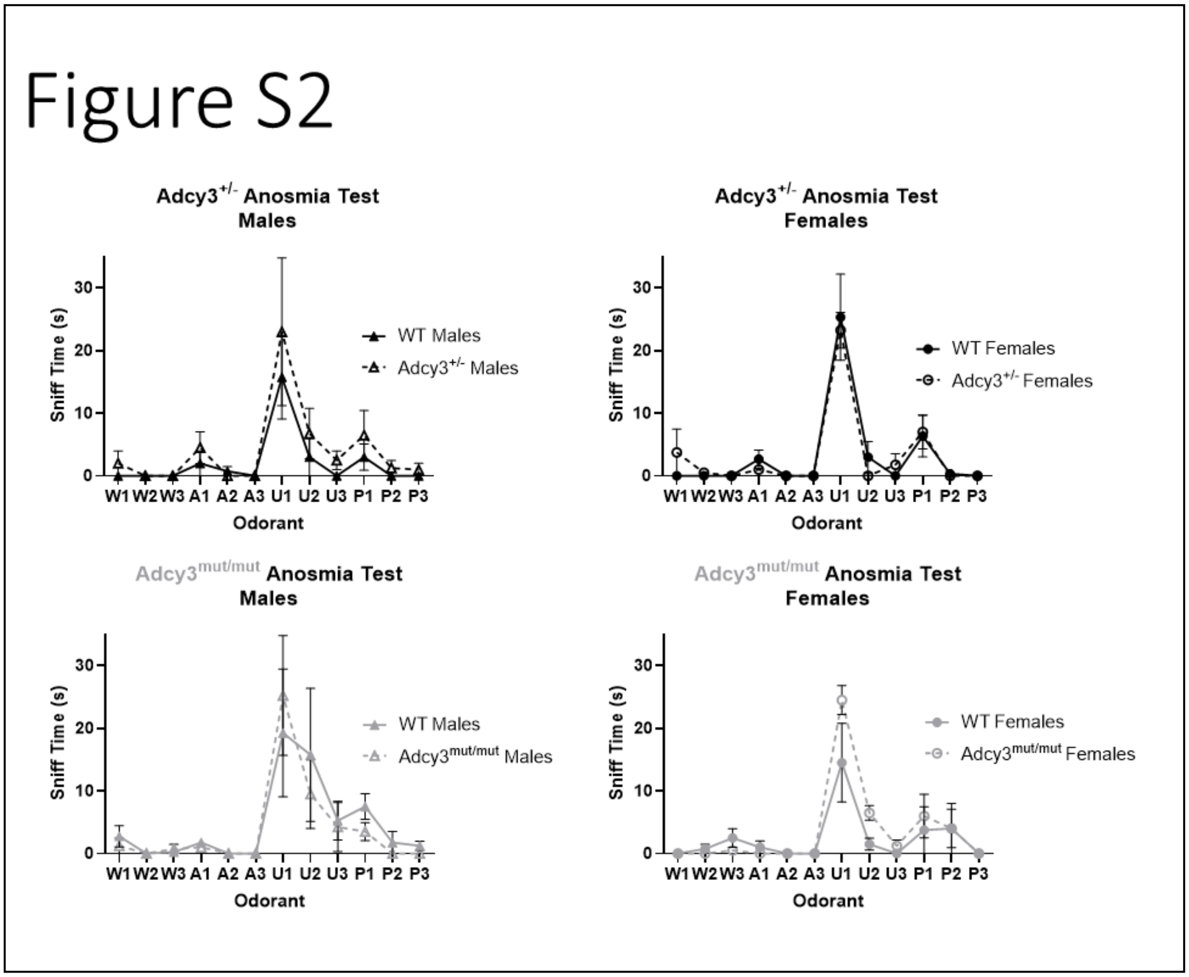

**Figure S3. Lean mass, non-adipose tissue weights at sac, and glucose area-under-the-curve (AUC) in Adcy3$^{+/-}$ and Adcy3$^{mut/mut}$.** No differences in lean mass prior to high-fat diet (HFD) start or after 8 weeks on diet (WOD). Among non-adipose tissues, there were no genotypic differences in weight, except that Adcy3$^{+/-}$ females (F) had slightly heavier pancreases than wild-type (WT) F. No differences in glucose AUC. Mean ± SEM. T-test, *p<0.05

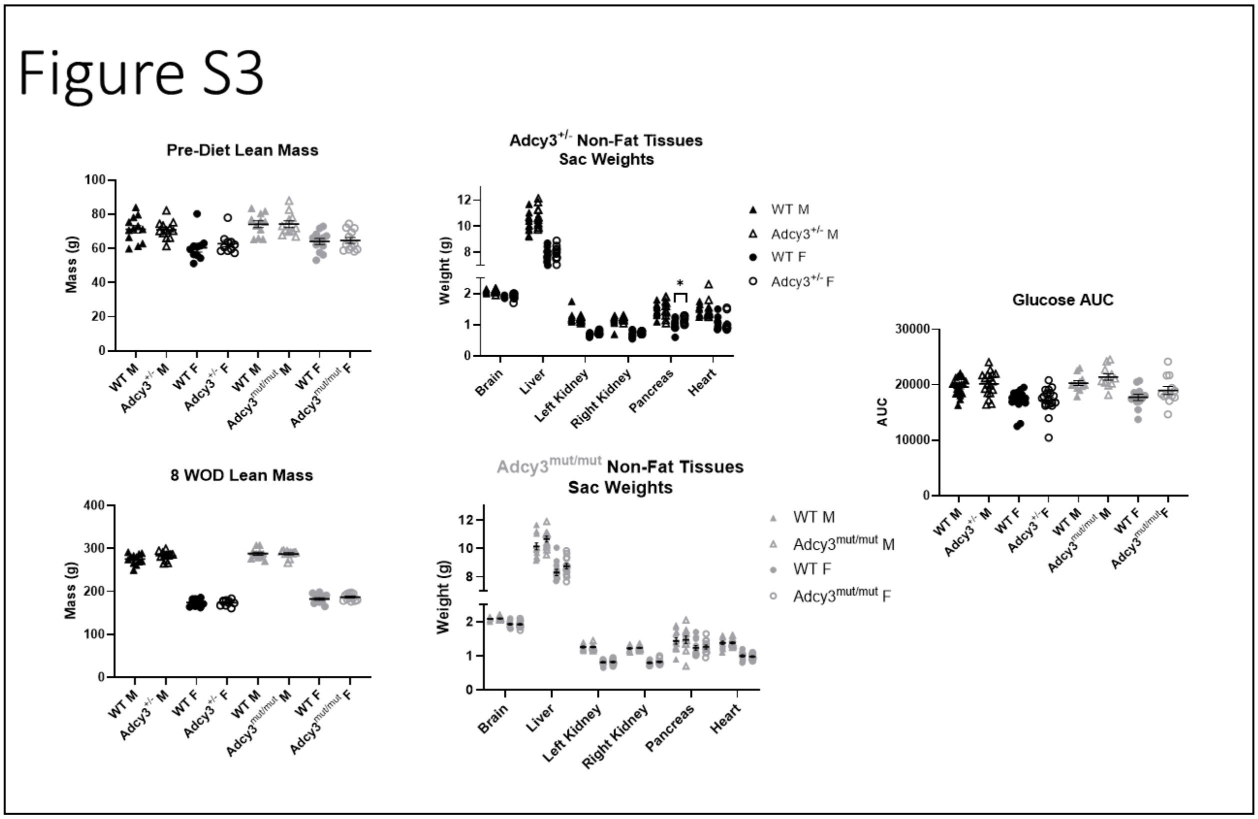

**Figure S4. Water intake and locomotor activity in the TSE PhenoMaster chambers in Adcy3$^{+/-}$ and Adcy3$^{mut/mut}$.** No differences in water intake or locomotor activity in any group. Mean ± SEM. ANCOVA.

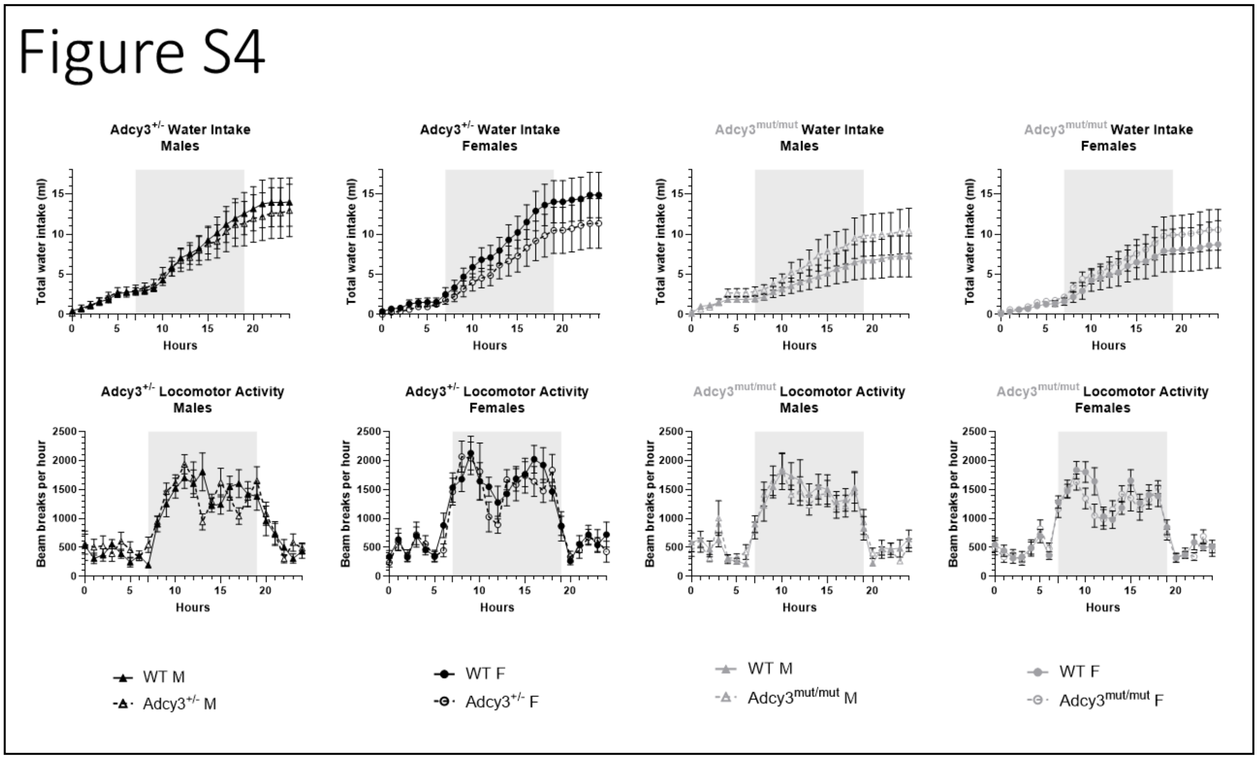

**Figure S5. Day 1 behavior in the novel object recognition test (NOR) in Adcy3$^{mut/mut}$.**

Objects A1 and A2 on Day 1 are identical. A1 is placed on the back side of the cage while A2 is on the front side. All groups spend roughly 50% of the time with both identical objects on Day 1. Wild-type (WT) males (M) do spend significantly less time with the object presented at the front of the cage. Mean ± SEM. T-test, **p<0.01

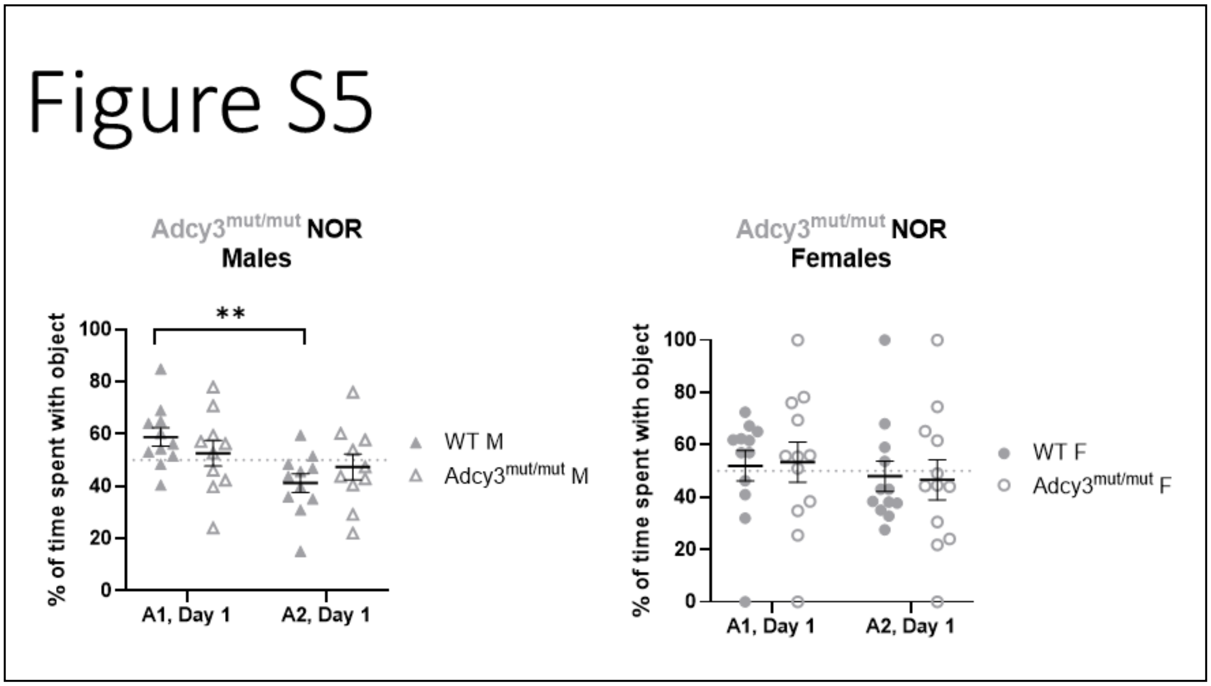

### Supplemental Tables

**Table S1. Primer sequences used in genotyping and RT-qPCR.**

| Primer | Forward Sequence | Reverse Sequence |
|---|---|---|
| *Adcy3* (genotyping) | TGTGCGGAACTCTGGGTCCT | AGATCTGGGCTGTGATGAGCAG |
| *Adcy3* (RT-qPCR) | AGAAGACCAAGACCGGAGTG | CATCTAGGTAGTCGCAGCGA |
| *β-actin* (RT-qPCR) | TGAGGTAGTCTGTCAGGTCCCG | ACCACTGGCATTGTGATGGACT |

**Table S2. Mean ± SEM in each behavior test outcome variable for each genotype-sex group.** *p<0.05, #0.05< p <0.10. P-values from student's t-tests conducted after separating by sex are depicted.

| Behavior | WT M | Adcy3$^{+/-}$ M | WT F | Adcy3$^{+/-}$ F | WT M | Adcy3$^{mut/mut}$ M | WT F | Adcy3$^{mut/mut}$ F |
|---|---|---|---|---|---|---|---|---|
| **OFT: Center Time (s)** | 155.78 ± 25.42 | 158.80 ± 27.36 | 62.09 ± 7.96 | 91.42 ± 12.80 | 111.40 ± 32.35 | 63.27 ± 8.03 | *54.18 ± 6.59 | *33.10 ± 5.09 |
| **OFT: Rearings** | 6.11 ± 1.23 | 7.90 ± 1.72 | 10.00 ±1.24 | 9.00 ± 1.58 | *3.20 ± 0.74 | *5.64 ± 0.88 | *9.42 ± 0.93 | *6.36 ± 0.79 |
| **OFT: Grooming Episodes** | 2.67 ± 0.62 | 2.60 ± 0.75 | 2.09 ± 0.53 | 2.08 ± 0.53 | #1.80 ± 0.73 | #4.36 ± 1.11 | 4.08 ± 0.65 | 3.00 ± 0.71 |
| **OFT: Line Crossings** | 15.44 ± 3.62 | 19.60 ± 3.53 | 31.27 ± 3.66 | 31.92 ± 2.91 | 8.20 ± 1.84 | 10.91 ± 2.04 | 27.17 ± 1.72 | 22.27 ± 3.47 |
| **OFT: Latency to leave center (s)** | 11.44 ± 3.94 | 15.10 ± 6.55 | 12.73 ± 3.48 | 7.00 ± 1.49 | 20.50 ± 4.36 | 18.36 ± 4.19 | 11.33 ± 2.59 | 8.45 ± 1.98 |
| **OFT: Fecal Boli** | 0.00 ± 0.00 | 0.00 ± 0.00 | 0.00 ± 0.00 | 0.00 ± 0.00 | 0.33 ± 0.17 | 0.20 ± 0.13 | 0.00 ± 0.00 | 0.00 ± 0.00 |
| **NOR: Day 1, A1 (%)** | 54.28 ± 3.22 | 54.18 ± 2.50 | 49.77 ± 1.84 | 55.92 ± 3.47 | *58.83 ± 3.58 | 52.65 ± 4.92 | 51.97 ± 5.79 | 53.35 ± 7.68 |
| **NOR: Day 1, A2 (%)** | 45.72 ± 3.22 | 45.82 ± 2.50 | 50.23 ± 1.84 | 44.08 ± 3.47 | *41.17 ± 3.58 | 47.35 ± 4.92 | 48.03 ± 5.79 | 46.65 ± 7.68 |
| **NOR: Day A2, (%)** | 38.52 ± 2.95 | 39.64 ± 1.73 | 36.91 ± 1.68 | 36.02 ± 1.46 | #36.47 ± 2.63 | #42.80 ± 1.66 | 37.19 ± 2.84 | 37.86 ± 3.52 |
| **NOR: Day 2, B, Novel (%)** | 61.48 ± 2.95 | 60.36 ± 1.73 | 63.09 ± 1.68 | 63.98 ± 1.46 | #63.53 ± 2.63 | #57.20 ± 1.66 | 62.81 ± 2.84 | 62.14 ± 3.52 |
| **FST: Mobile Counts** | 20.10 ± 2.83 | 16.92 ± 3.01 | 28.60 ± 3.01 | 25.00 ± 2.08 | *23.70 ± 3.72 | *7.44 ± 1.20 | 21.45 ± 3.31 | 14.80 ± 2.39 |
| **FST: Immobile Counts** | 41.33 ± 2.83 | 43.08 ± 3.01 | 31.40 ± 3.01 | 35.00 ± 2.08 | *36.30 ± 3.72 | *52.56 ± 1.20 | 38.55 ± 3.31 | 45.20 ± 2.39 |
